# *Mycobacterium tuberculosis* infection associated immune perturbations correlate with antiretroviral immunity

**DOI:** 10.1101/2023.07.14.548872

**Authors:** Burcu Tepekule, Lisa Jörimann, Corinne D. Schenkel, Lennart Opitz, Jasmin Tschumi, Rebekka Wolfensberger, Kathrin Neumann, Katharina Kusejko, Marius Zeeb, Lucas Boeck, Marisa Kälin, Julia Notter, Hansjakob Furrer, Matthias Hoffmann, Hans H. Hirsch, Alexandra Calmy, Matthias Cavassini, Niklaus D. Labhardt, Enos Bernasconi, Karin J Metzner, Dominique L. Braun, Huldrych F. Günthard, Roger D. Kouyos, Fergal Duffy, Johannes Nemeth, the Swiss HIV Cohort Study

## Abstract

Infection with Mycobacterium tuberculosis (MTB) remains one of the most important opportunistic infections in people with HIV-1 (PWH). While active Tuberculosis (TB) leads to rapid progression of immunodeficiency in PWH, the interaction between MTB and HIV-1 during the asymptomatic phase of both infections remains poorly understood.

In a cohort of individuals with HIV (PWH) with and without suppressed HIV-1 viral load, the transcriptomic profiles of peripheral blood mononuclear cells (PBMC) clustered in individuals infected with Mycobacterium tuberculosis (MTB) compared to carefully matched controls. Subsequent functional annotation analysis disclosed alterations in the IL-6, TNF, and KRAS pathways. Notably, MTB-associated genes demonstrated an inverse correlation with HIV-1 viremia, evident at both on individual gene level and when employed as a gene score.

In sum, our data show that MTB infection in PWH is associated with a shift in the activation state of the immune system, displaying an inverse relationship with HIV-1 viral load. These results could provide an explanation for the observed increased antiretroviral control associated with MTB infection in PWH.

## Introduction

Globally, HIV-1 remains one of the most important risk factors for the development of active Tuberculosis (TB) [1,2]. Individuals with HIV-1 and active TB suffer from earlier TB dissemination, increased risk of faster progression and TB drug resistance[1,3].

However, active TB represents only the most severe manifestation within the spectrum of Mycobacterium tuberculosis (MTB) infection. There are ample efforts to advance the field beyond the binary concept of infection versus disease into a more granular description of the disease spectrum[4]. While the exact numbers remain uncertain, it is well accepted that less than 10% of MTB-infected HIV-uninfected individuals ever develop active disease[2]. Similarly, only a minority of people with HIV-1 (PWH) exposed to MTB will progress to active TB [1,5].

Until now, the sole method of verifying exposure to MTB in clinical practice in the absence of symptomatic TB has been through the identification of MTB specific T cells either by stimulating T cells intradermally (tuberculin skin test) or circulation, known as Interferon Gamma Release Assays (IGRA). However, this approach has significant limitations. The presence of MTB specific T cells could indicate one of three possibilities. Firstly, it could suggest the presence of bacteria that are in a state of reversible latency and have the potential to cause disease. Secondly, it could indicate bacteria that are in a state of practically “irreversible” latency, requiring an excessive and unsustainable amount of immunosuppression to become active again, incompatible with the host’s survival. Lastly, it could signify a memory T cell response following the eradication of the infection [6]. Additionally, the efficacy of IGRA relies on the functionality of T cells, which can be impaired due to the replication of HIV-1[1].

In this work, we hypothesized that exposure to MTB does not only induce an antigen specific T cell response but also impacts the activation state of the innate immune system analogous to the trained immunity phenotype described after BCG vaccination [7]. This hypothesis originated from data we generated in an *in vivo* model of asymptomatic MTB infection, and from heterologous effects observed in a large clinical cohort [8–11]. In people with HIV-1, MTB infection but not active TB was associated with significantly reduced HIV-1 viral set point and with a lower frequency of common opportunistic infections[10]. Consequently, we investigated the immunological modifications induced by MTB infection and their relationship with the HIV-1 viral set point, considering the significant and consistent impacts observed at the population level[10].

## Material and Methods

### Selection and matching of study population

We sourced peripheral blood mononuclear cells (PBMC) from 36 people living with HIV (PWH) participating in the Swiss HIV Cohort Study (SHCS). The SHCS is a prospective, longitudinal, multicentre, representative cohort study that collects demographic and clinical data from people with HIV-1 twice a year and stores plasma and PBMC samples in a biobank[12]. MTB positivity was defined based on one or more positive tuberculosis (TB) tests (IGRA, TST, or both) in the absence of diagnosed pulmonary or extrapulmonary TB. We selected only those with unequivocal TB test results. These samples were then categorized into four groups based on their *Mycobacterium tuberculosis* infection (MTB) and HIV infection status: *i)* MTB positive and viremic (MTB+ HIV^vir^; n=7), *ii)* MTB positive and suppressed (MTB+ HIV^supp^; n=11), *iii)* MTB negative and viremic (MTB-HIV^vir^; n=7), *iv)* MTB negative and suppressed (MTB-HIV^supp^; n=11). For each sample, we used the TB test closest to the time of blood collection, with 78% of samples coinciding with the TB testing date. PWH were classified as suppressed if their HIV viral load was below 500 copies/ml and as viremic if it was above or equal to this threshold. As outlined in Table 1, all patients categorized “supp” and detected by the algorithm had undetectable HIV-1 viral load. We included only samples collected post-2000 to minimize the risk of cell degradation. Our selection was also influenced by practical constraints, such as the availability of sample aliquots and the viability of PBMCs. Individuals with other opportunistic infections were excluded (detailed list in Supplementary Table 2).

**Table 1.** Characteristics of suppressed PWH. Shown are median and interquartile range or % for MTB+ HIV^supp^ (n=11), MTB-HIV^supp^ (n=11) and all HIV^supp^ (n=22) with corresponding p-values from two sample t-tests. TST= Tuberculin skin test, IGRA= Interferon gamma release assay.

|  |  | MTB+ HIV<sup>supp</sup> | MTB- HIV<sup>supp</sup> | all HIV<sup>supp</sup> | p-value |
| --- | --- | --- | --- | --- | --- |
|  |  | 11 | 11 | 22 |  |
| CD4 |  | 480 | 482 | 481 |  |
| (median [IQR]) |  | [458, 742] | [404, 749] | [425, 839] | 0.948 |
| Viral load |  | 0 | 0 | 0 |  |
| (median [IQR]) |  | [0, 0] | [0, 0] | [0, 0] |  |
| TB test type (%) | TST | 2 (18.2) | 3 (27.3) | 5 (22.7) | 0.631 |
|  | IGRA | 9 (81.8) | 8 (72.7) | 17 (77.3) |  |
| Sex (%) | Male | 9 (81.8) | 9 (81.8) | 18 (81.8) | 1 |
|  | Female | 2 (18.2) | 2 (18.2) | 4 (18.2) |  |
| Ethnicity (%) | White | 9 (81.8) | 9 (81.8) | 18 (81.8) | 1 |
|  | Black | 1 ( 9.1) | 1 ( 9.1) | 2 ( 9.1) |  |
|  | Hispano-american | 1 ( 9.1) | 1 ( 9.1) | 2 ( 9.1) |  |

**Table 2.** Characteristics of viremic PWH. Shown are median and interquartile range or % for MTB+ HIV^vir^ (n=7), MTB-HIV^vir^ (n=7) and all HIV^vir^ (n=14) with corresponding p-values from two sample t-tests. IGRA= Interferon gamma release assay.

|  |  | MTB+ HIV<sup>vir</sup> | MTB- HIV<sup>vir</sup> | all HIV<sup>vir</sup> | p-value |
| --- | --- | --- | --- | --- | --- |
|  |  | 7 | 7 | 14 |  |
| CD4 |  | 453 | 453 | 453 |  |
| (median [IQR]) |  | [373, 656] | [373, 653] | [355, 677] | 0.93 |
| Viral load |  | 33'643 | 120'000 | 33'872 |  |
| (median [IQR]) |  | [18'461, 93'725] | [21'782, 700'000] | [18'343, 472'678] | 0.258 |
| TB test type (%) | IGRA | 7 (100.0) | 7 (100.0) | 14 (100.0) |  |
| Sex (%) | Male | 6 (85.7) | 6 (85.7) | 12 (85.7) | 1 |
|  | Female | 1 (14.3) | 1 (14.3) | 2 (14.3) |  |
| Ethnicity (%) | White | 7 (100.0) | 7 (100.0) | 7 (100.0) |  |

We considered sex, ethnicity, and CD4 cell counts when matching MTB- and MTB+ individuals within each HIV status group. Matching process involves first identifying all potential matches with the same sex and ethnicity, then selecting the individual with the closest CD4 cell count. Once a pair was matched, they were removed from the pool to prevent duplication in the matching process (Supplementary Figure 2).

### Flow Cytometric Analysis

For flow cytometric analysis, cells underwent two washes with PBS to eliminate any remaining stimulation compounds. Surface proteins were then stained at 4°C for 30 minutes using the following markers: NIR Zombie dye APC-Cy7 (Biolegend, Cat No. 423105), CD20 PE (Biolegend, Cat No. 302306), CD3 AF700 (Biolegend, Cat No. 300324), CD69 FITC (Biolegend, Cat No. 310904), CD56 BV510 (Biolegend, Cat No. 304608), CD16 PE-Dazzle (Biolegend, Cat No. 302054), HLA-DR BV605 (Biolegend, Cat No. 307640), CD11b PE-Cy7 (Biolegend, Cat No. 301322), and CD14 BV421 (Biolegend, Cat No. 301829). Post-staining, the cells were washed again with PBS and fixed using Cytofix/Cytoperm buffer (BD Biosciences, Cat No. 554722) for 20 minutes at 4°C. Following another wash, cells were resuspended in PBS. The flow cytometry was conducted on an LSR II Fortessa 4L equipped with a High throughput sampler (HTS) (Supplementary Figure 3). Data analysis was performed using FlowJo version 10.0.8r1 and R software.

### RNA Isolation and Sequencing

Following cell stimulation, the samples were centrifuged for 5 minutes at 600 g. RNA isolation was then performed using the RNeasy Micro Kit (Qiagen), following the manufacturer’s instructions. This process was carried out either manually or utilizing the QIAcube (Qiagen). The quantity of isolated RNA was assessed on a TapeStation (Agilent) employing a high-sensitivity RNA ScreenTape. For RNA sequencing (RNAseq), 20 ng of total RNA was used as input. We utilized the SmartSeq2 library preparation method, and sequencing was conducted with a 1×100 bp single-end read configuration (Illumina). Each sample was sequenced to a minimum depth of 30 million reads using the NovaSeq6000 system (Illumina), according to the manufacturer’s guidelines. All sequencing was performed at the Functional Genomics Centre Zurich (FGCZ).

### RNAseq analysis

The RNA sequencing raw reads were first cleaned by removing adapter sequences, trimming low quality ends, and filtering reads with low quality (phred quality <20) using Fastp (Version 0.20). Sequence pseudo alignment of the resulting high-quality reads to the human reference genome (build GRCh38.p13) and quantification of gene level expression (gene models based on GENCODE release 37) was carried out using Kallisto (Version 0.46.1)[13]. Differential expression was computed using the generalized linear model as implemented in the Bioconductor package DESeq2 (R version: 4.2.0, DESeq2 version: 1.36.0)[14]. Genes showing altered expression with adjusted (Benjamini and Hochberg method) p-value <0.05 were considered differentially expressed. Batch effects were corrected using the removeBatchEffect function from the limma package (v3.54.1)[15]. Due to technical issues, certain samples were not successfully sequenced. These samples included 2 matched MTB-/+ pairs (4 samples) belonging to HIV^supp^, 2 samples belonging to MTB-HIV^vir^, and 1 sample belonging to MTB+ HIV^vir^ from the untreated pool (negative control). Consequently, this lowered the number of matched samples from 11 to 9 for HIV^supp^, and led to an unmatched HIV^vir^ population with 5 MTB- and 6 MTB+ samples. Given the already low number of samples available, for the dimensionality reduction and clustering analysis, we proceeded with the unmatched HIV^vir^ population since the MTB-/+ is a binary categorization. However, this mismatch was a bigger concern when calculating the viral load correlations. Therefore, for every sample that failed sequencing, its corresponding matched sample was also excluded, resulting in a final count of 4 matched pairs for HIV^vir^. Dimensionality reduction and clustering analysis is performed on this matched population as well, given in Supplementary Figure 4.

### Dimensionality reduction and clustering analysis

K-means clustering was applied to normalized gene counts using the k-means function in the R stats package. We focused on genes with a differential expression p-value below 0.01. The number of clusters was set to four, corresponding to the four sample groups for each HIV phenotype (unstimulated MTB+, stimulated MTB-, stimulated MTB+ with PMA/Ionomycin, and stimulated MTB-with PMA/Ionomycin). For our Gene Set Enrichment Analysis (GSEA), we utilized the Hallmark database [ref]. When visualizing the heatmaps, z-score normalization was applied to the log2-transformed normalized read counts for each gene.

### Correlation calculations between viral load and gene expression values

We analyzed the correlation between log10-transformed viral RNA counts and log10-transformed FPKM values, focusing on conserved and non-conserved transcripts with criteria of abs(LogFC)>1 and p-value<0.01. The distributions of these correlation coefficients were visualized in boxplots, and a one sided t-test was used to determine whether the non-conserved transcripts had a significantly greater correlation value.

Permutation tests were conducted by creating 1,000 random subsets, each matching the size of our conserved transcript gene pool, drawn from the entire transcript set. For each subset, average FPKM values were computed, taking into account the direction of gene regulation. These values were then correlated with log10-transformed viral RNA counts to assess the significance of our observed correlations for the conserved transcripts.

### Integration and comparison to other datasets

To validate our results, we utilized an independent dataset: a cohort comprising PWH with either MTB infection or active TB (PWH MTB+, PWH developing active TB)[16]. The dataset from Kaforou *et al*. provided background-subtracted, quantile-normalized whole blood microarray expression data (downloaded from NCBI GEO, GSE37250). We applied a log2 transformation to this data. Signature scores [17] were calculated per patient by subtracting the expression level of the downregulated genes from the upregulated genes in MTB+ compared to MTB-, given the direction of regulation determined in our study. We applied one-tailed t-tests to compare the distribution of signature scores statistically.

#### MTB infection perturbs the transcriptome of PWH with suppressed viral load

To investigate the impact of MTB infection on the transcriptome of people with HIV (PWH), we conducted a detailed analysis using peripheral blood mononuclear cells (PBMC). We focused PWH with a suppressed viral load to adjust for the potential effects of a replicating HIV-1 infection on the transcriptome. We carefully matched 22 PWH in pairs based on their CD4 cell count, ethnicity, and sex to minimize further confounding variables (Table 1).

Our analysis revealed distinct transcriptome profiles between HIV-suppressed individuals with and without MTB infection (MTB+ HIV^supp^ and MTB-HIV^supp^), particularly in non-activated PBMCs, as shown by dimensionality reduction and clustering (Figure 1A).

**Figure 1.**
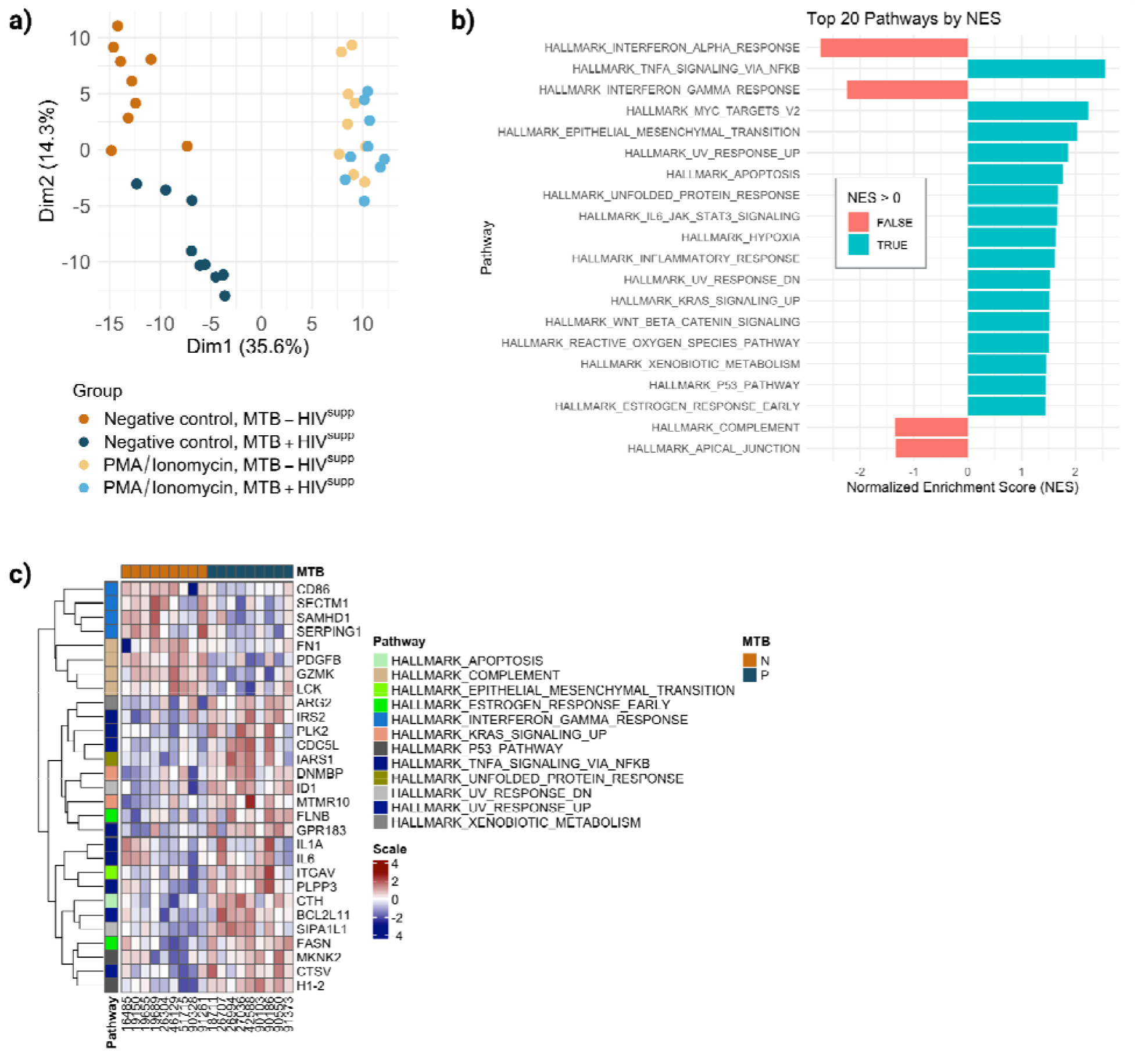
MTB Perturbations in suppressed PWH. **a)** Dimensionality reduction and clustering analysi for HIV-suppressed individuals with and without MTB infection (MTB+ HIV^supp^ and MTB-HIV^supp^). Gene with a p-value lower than 0.01 are used for the analysis, corresponding to 293 genes out of the 21493 analyzed. In MTB+ patients compared to MTB-patients, 126 out of 293 genes were upregulated and 167 were downregulated. **b)** First 20 pathways with the highest normalized enrichment score (NES) based on the gene-set enrichment analysis (GSEA). **c)** Heatmap displaying the leading edge genes of these pathways filtered by p-value <0.01.

**Figure 2.**
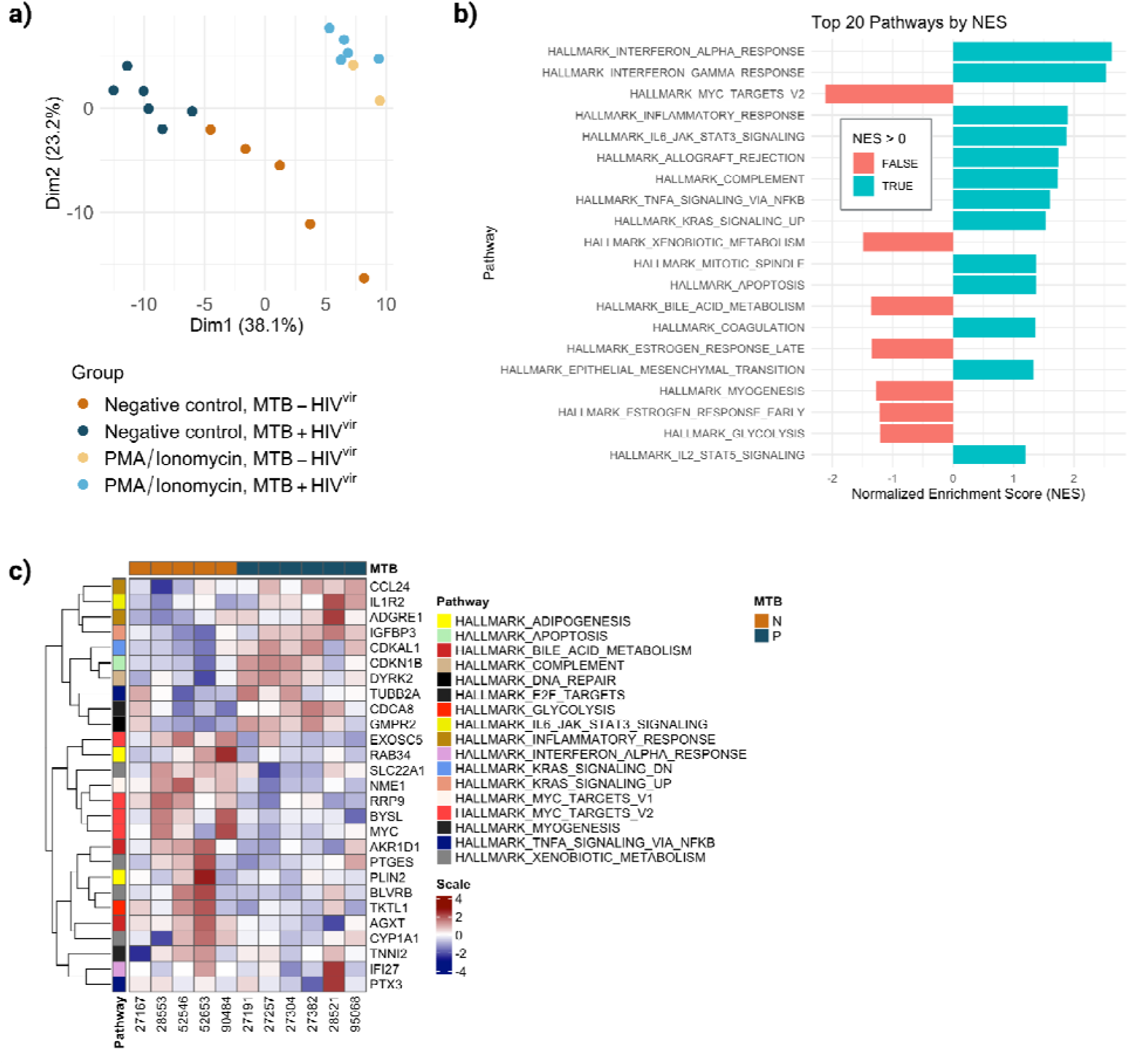
MTB Perturbations in viremic PWH. **a)** Dimensionality reduction and clustering analysis for HIV-viremic individuals with and without MTB infection (MTB+ HIV^vir^ and MTB-HIV^vir^). Genes with a p-value lower than 0.01 are used for the analysis, corresponding to 166 genes out of the 21467 analyzed. In MTB+ patients compared to MTB-patients, 68 out of 166 genes were upregulated and 98 were downregulated. **b)** First 20 pathways with the highest normalized enrichment score (NES) based on the gene-set enrichment analysis (GSEA). **c)** Heatmap displaying the leading edge genes of these pathways filtered by p-value <0.01.

Using gene-set enrichment analysis (GSEA), we ranked the pathways with the highest normalized enrichment score (NES) (Figure 1B) and displayed the leading edge genes of these pathways in a heatmap (Figure 1C). Flow cytometry analysis revealed no significant differences in the main PBMC subpopulations, suggesting that the transcriptomic changes are not due to variations in cell type frequencies (Supplementary Figure 1a and b).

#### MTB infection perturbs the transcriptome of PWH with replicating HIV-1 virus

Next, we examined the impact of MTB infection on HIV viral load in viremic PWH. We analyzed 14 viremic PWH, both with and without MTB co-infection (MTB+ HIV^vir^ and MTB-HIV^vir^), matched for CD4 cell count, ethnicity, and sex (Supplementary Figure 2) (Table 2).

Consistent with our findings in suppressed PWH, dimensionality reduction and clustering analysis identified distinct transcriptomic profiles between MTB+ HIV^vir^ and MTB-HIV^vir^ (Figure 2A). The clear separation between the MTB+ and MTB-groups emphasizes the significant impact of MTB infection on the transcriptome, an effect that remained notably detectable despite the extensive perturbations caused by HIV-1 infection[18].

We then investigated the transcriptional changes due to MTB infection in viremic PWH with GSEA and ranked the pathways with the highest NES (Figure 2B), and displayed the leading edge genes of these pathways in a heatmap (Figure 2C).

In sum, in subjects with MTB infection, a noteworthy and sustained upregulation was discerned in three hallmark pathways when compared to their non-infected counterparts. These pathways include tumor necrosis factor (TNF) signaling, KRAS signaling, and interleukin-6 (IL-6) signaling, irrespective of the occurrence of human immunodeficiency virus type 1 (HIV-1) viremia, as illustrated in Figure 1b and Figure 2b.

#### Testing the MTB - HIV-1 interaction on single gene level

At the level of molecular pathways, our findings indicate that MTB infection in PWH is linked to increased activation of pro-inflammatory pathways, regardless of the presence of HIV-1 viremia. Given the observation that MTB infection is correlated with a reduction in HIV-1 viral load, we hypothesized that the heightened activity of these upregulated pathways may play a role in explaining this observed phenomenon.

To test this association on a single gene level, we identified ‘conserved transcripts’ as genes that were consistently differentially expressed in the same direction in MTB+ individuals compared to MTB-irrespective of HIV viremia status (full list in Supplementary Table 1). This categorization of genes identifies genes associated with MTB infection as a common denominator across HIV-1 viremic and HIV-1 suppressed patients. We were unable to identify any enriched pathways associated with these genes. We examined the correlation between HIV-1 viral load and the expression levels of these genes across different fold-change thresholds (Figure 3A). We adjusted these correlations to account for the direction of gene regulation. For instance, if a gene is downregulated in MTB+ compared to MTB-, but its expression is inversely correlated with viral load, the correlation value would be inverted (multiplied by -1). Notably, as we raised the fold-change threshold — thereby focusing on genes with greater differences in expression according to the MTB status — we found a stronger and more statistically significant negative correlation with viral load, speaking to the strength of the effect.

To determine the probability of these correlations arising by chance, we employed a permutation test. This involved generating multiple random subsets from the pool of all transcripts and calculating the average correlation between the expression levels and the viral load in these randomized sets. We then compared this distribution to the correlations observed in our conserved transcripts. The results of the permutation test gave a p-value of less than 2.2e-16 (Supplementary Figure 3), demonstrating that the negative correlation between the expression of conserved transcripts and viral load was highly unlikely to occur by chance.

To understand which cells are most actively engaged in this mitigating effect, we analyzed the proportions of different immune cells expressing these transcripts (Figure 3b). Figure 3b illustrates that fifty percent of the genes were linked to cells originating from the innate immune system, while twenty-five percent were associated with the adaptive immune system.

### Testing the MTB-HIV-1 interaction integrating the single gene information into a gene score

To quantify the collective expression pattern of the conserved transcripts defined above, we calculated the signature score [17]using the 32 genes provided in Supplementary Table 1. We first compared this score between viremic and suppressed patients, suggesting a significant difference between the score values (Figure 4a). We then calculated the correlation of this signature score with the HIV-1 viral load in PWH, suggesting a correlation coefficient of -0.31 (Figure 4B).

Previous studies have shown that active tuberculosis (TB) is associated with increased HIV-1 viral load [1,10,19]. We therefore hypothesized that increased antiretroviral control observed in asymptomatic MTB infection might be lost during active TB. To test this hypothesis, we focused on the “conserved transcripts” that are upregulated during asymptomatic MTB infection and inversely associated with HIV-1 viral set point, predicting these would be downregulated in active TB. We analyzed their expression in an independent cohort of PWH with asymptomatic MTB or active TB infection from Malawi and South Africa[16]. Our analysis showed that PWH developing active TB had a significantly lower signature compared to PWH MTB+ when calculated over these gene sets (Figure 5). These preliminary findings support our hypothesis that losing control over MTB infection also affects key genes associated with retroviral control.

**Figure 3.**
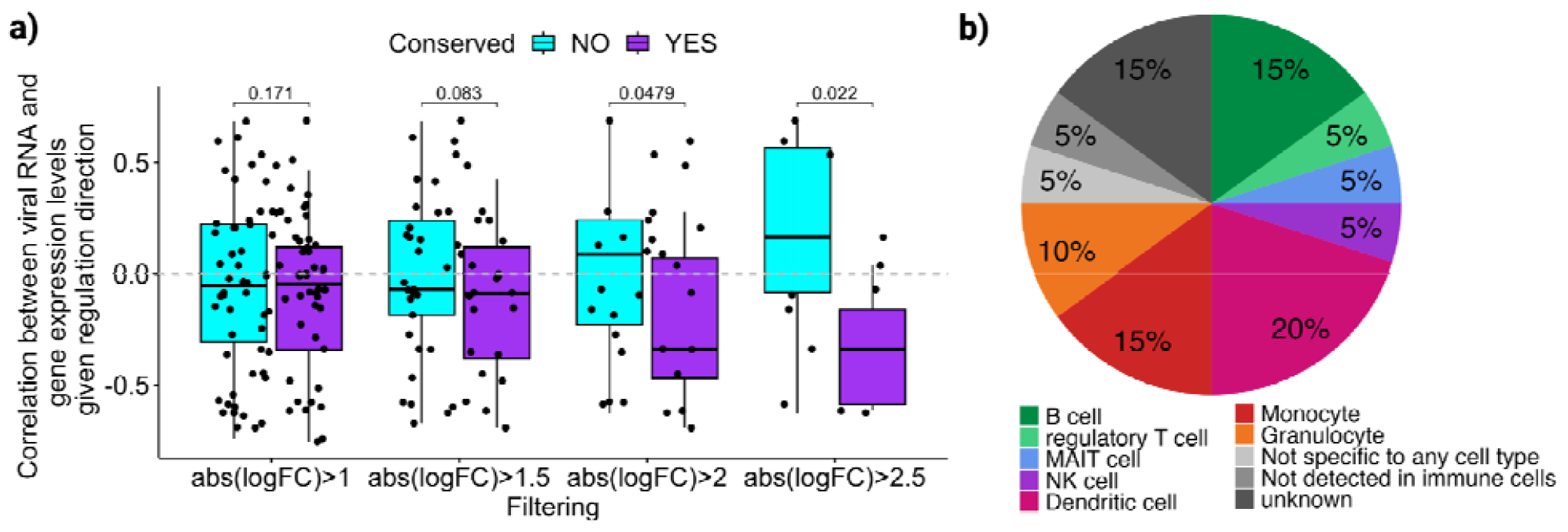
Correlation between HIV-1 viral load and the expression levels of conserved transcripts. **a)** Correlation between HIV-1 viral load and the expression levels of carried over transcripts acros different fold-change thresholds. **b)** Percentages of different immune cells expressing the conserved transcripts with a negative correlation to viral load (18 transcripts out of 32), according to the human protein atlas[32].

**Figure 4.**
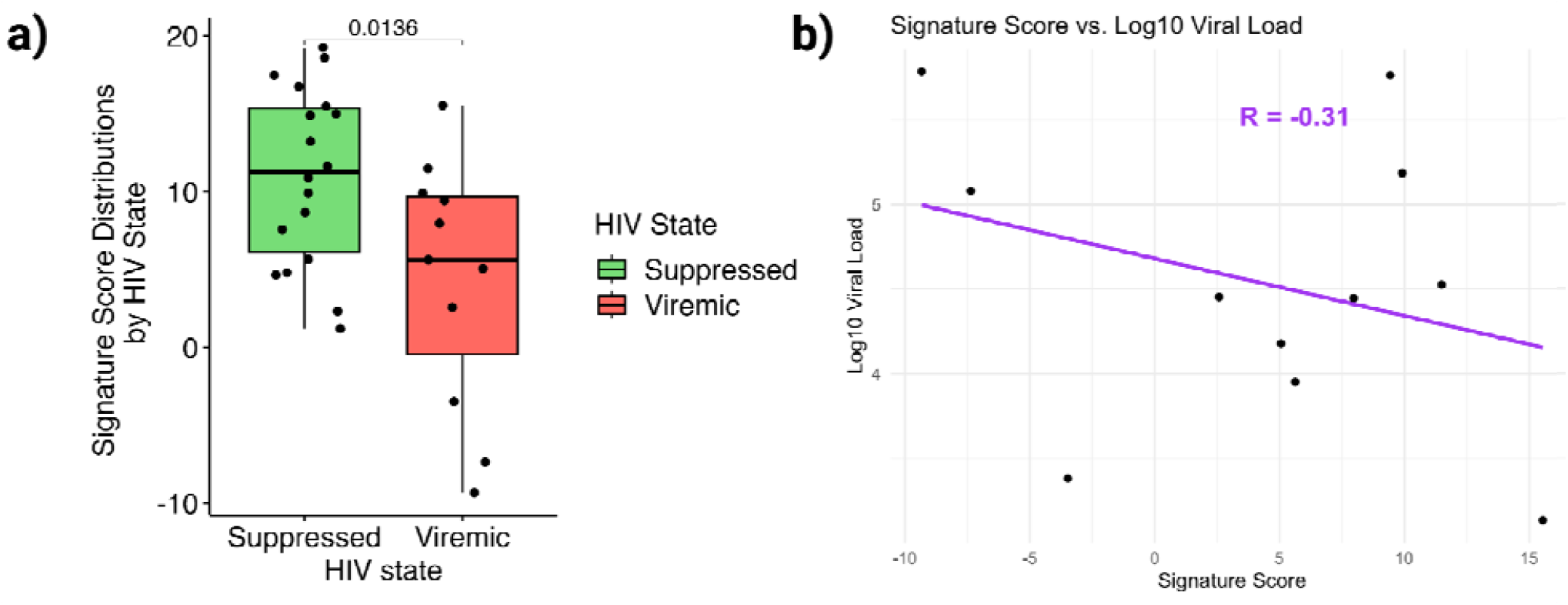
**a)** Signature score distributions of all carried over transcripts by HIV state. **b)** Correlation between HIV-1 viral load and the signature score of carried over transcripts in PWH.

**Figure 5.**
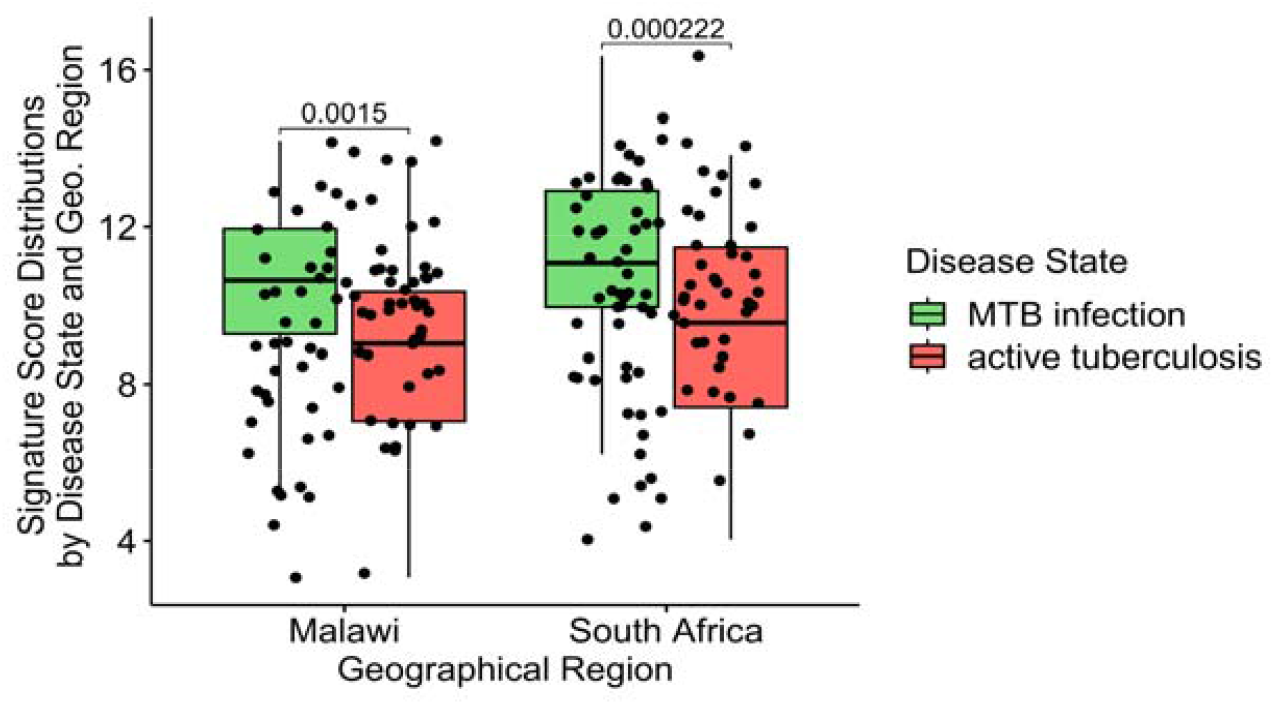
Signature Score Distributions in active TB. Signature score distributions including all 9 carried over transcripts that were present in our study and in the African cohort.

## Discussion

In this study, we have shown that MTB infection correlates with transcriptional changes circulating PBMCs in PWH, independently of HIV-1 viral load. The MTB-associated transcriptional changes were inversely correlated with HIV-1 viremia, suggesting a potential causal link between MTB infection and improved control of HIV-1 replication.

The consistent upregulation of TNF alpha signaling via NF-_κ_B and the IL6-JAK-STAT3 pathway in MTB infection irrespective of HIV-1 viremia suggest a sustained inflammatory response. The pro-inflammatory cytokines IL-6 and TNF-a have traditionally been associated with active disease in various cross-sectional studies comparing active TB patients to treated patients or healthy controls[20,21]. Also, IL-6 levels have been shown to be predictors of unfavorable outcomes in patients with active TB[22]. Conversely, IL-6 signaling has been hypothesized to be advantageous by inhibiting detrimental Type 1 Interferon responses[23]. An independent study found increased IL-6 and TNF-a levels on a protein level to be associated with MTB infection in the absence of active TB, further inducing confidence in our findings[24]. Likewise, TNF signaling has been linked to disease progression of both MTB and HIV-1 infection[25–29]. Our investigation adds a different aspect to these pro-inflammatory signaling during asymptomatic MTB infection: we demonstrate the activation of these very pathways in PWH who will remain free from active TB for years to decades[5].

Subsequently, our objective was to establish a correlation between alterations in the inflammatory state and potential heterologous effects, specifically focusing on HIV-1 control[10]. We systematically examined these perturbations on the single gene and gene score levels, unveiling an inverse relationship between the expression of MTB infection associated genes and HIV-1 control. It is noteworthy that our investigation contributes to the existing body of literature by emphasizing that these genes were defined by MTB infection status (and not, like in other investigations, by HIV-1 phenotype (for example[30])), therefore implying a heterologous mechanism. Also, it is important to note that these genes were not enriched for a specific pathway. Thus, it is important to recognize that the genes we identified are not the sole determinants of HIV control, but rather exemplify the concept that certain features of the innate immune response induced by an asymptomatic MTB infection are consistent across various HIV viremic states. While our focus on these conserved transcripts has provided surprising inverse correlations, they represent just one aspect of the complex interplay between MTB infection and HIV control. Future studies are needed to unravel the full spectrum of immune responses and mechanisms contributing the control of HIV replication.

Our findings suggest that transcriptional changes linked to enhanced antiretroviral immunity in MTB-infected individuals are reversed during active TB. In a study involving PWH from Malawi and South Africa, we evaluated the expression of genes upregulated during asymptomatic MTB infection and inversely correlated with HIV-1 viral set point in our initial analysis. We observed a significant decrease in their expression among individuals with active TB compared to those with asymptomatic MTB infection, supporting the hypothesis of retroviral control loss during active TB.

Our study possesses notable strengths. Firstly, the Swiss HIV Cohort Study (SHCS) facilitates precise matching of individuals with and without Mycobacterium tuberculosis (MTB) infection, providing vital clinical insights into the absence of TB. Particularly in people with HIV (PWH), active TB may present sub-clinically[31], highlighting the importance of a highly granular, clinical phenotype for the definition of MTB infection. Secondly, the study benefits from extensive longitudinal follow-up, allowing the identification of individuals with MTB-specific T cell responses who have remained free of active tuberculosis over subsequent years to decades.

Our study has limitations: the cohort comprises only PWH from a Swiss longitudinal study, potentially introducing selection biases. Additionally, while bulk RNA sequencing provides a comprehensive view, it may miss subtleties detectable by single-cell sequencing. Furthermore, the study involves a limited number of patients, though meticulous patient matching enhances its robustness. Despite this limitation, a consistent signal emerges among MTB-infected patients, highlighting the potential significance of MTB infection even within a small cohort, warranting further investigation.

In aggregate we propose that, in specific individuals exposed to MTB, the heightened inflammation does not unequivocally translate into active TB. Also, our findings suggest that in a subset of patients, MTB associated transcriptional changes are correlated with enhanced control of HIV-1. Consequently, our data seamlessly align with the evolving idea of MTB infection as a spectrum of diseases, wherein stages of MTB infection differently influence the immune system.

## Funding information

This study was supported by the Swiss National Science Foundation (SNF) grant 310030_200407, the Theodor und Ida Herzog Stiftung, SHCS Project Funding 857 and the NIH Grant U19AI100627. The SHCS is funded by different sources, mainly by the Swiss National Science Foundation (SNSF), companies and the Swiss HIV Cohort Research Foundation. Since 2000, the Swiss National Science Foundation (SNSF) has been the major funding organization of the SHCS.

## Acknowledgements

We would like to thank all the SHCS study participants, doctors and nurses.

## Conflict of interest

A. C. received grants from Merck Sharp & Dohme (MSD), ViiV Healthcare, and Gilead Sciences for unrelated research. R. D. K. received grants from Gilead Sciences and National Institutes of Health (NIH) for unrelated research. D. L. B. received honoraria for working on the advisory board of Gilead Sciences, Merck, ViiV, Pfizer, and AstraZeneca. D. L. B. received honoraria for presentations from Gilead Sciences and Merck. E. B. received grants from MSD for unrelated research. E. B. received payments for travel reimbursement from ViiV, MSD, Gilead Sciences, Pfizer, and Abbvie. E. B. received honoraria for working on the advisory board of ViiV, MSD, Pfizer, Gilead Sciences, AstraZeneca, and Ely Lilly. H. H. H. received honoraria for working on the advisory board of AiCuris, Merck, Vera Dx, and Molecular Partners. H. H. H. received honoraria for presentations from Merck, Gilead Sciences, Biotest, and Vera Dx. J. N. received honoraria for presentations from Oxford Immunotec, Gilead and ViiV. H. F. G. received honoraria for working on the advisory board of Gilead Sciences, Merck, ViiV, Janssen, Johnson and Johnson, Novartis, and GlaxoSmithKline (GSK). H. F. G. received payments for travel reimbursements from Gilead Sciences. H. F. G. received grants from NIH, Yvonne Jacob Foundation, and Gilead Sciences. All other authors report no potential conflicts.

## Notes

### Summary of Updates

This version of the manuscript has been revised to update the results, figures and discussion.

## References

1. Kwan CK, Ernst JD. HIV and tuberculosis: a deadly human syndemic. Clinical microbiology reviews. 2011; 24(2):351–376.

2. Pai M, Behr MA, Dowdy D, et al. Tuberculosis. 2016; 2:16076–23.

3. Wells CD, Cegielski JP, Nelson LJ, et al. HIV infection and multidrug-resistant tuberculosis: the perfect storm. The Journal of infectious diseases. 2007; 196 Suppl 1(1):S86–107.

4. Esmail H, Macpherson L, Coussens AK, Houben RMGJ. Mind the gap – Managing tuberculosis across the disease spectrum. Ebiomedicine. 2022; 78:103928.

5. Zeeb M, Tepekule B, Kusejko K, et al. Understanding the Decline of Incident, Active Tuberculosis in People With Human Immunodeficiency Virus (HIV) in Switzerland. Clin Infect Dis. 2023;.

6. Emery JC, Richards AS, Dale KD, et al. Self-clearance of Mycobacterium tuberculosisinfection: implications for lifetime risk and population at-risk of tuberculosis disease. Proceedings of the Royal Society of London Series B: Biological Sciences. 2021; 288(1943):20201635–8.

7. Ziogas A, Bruno M, Meel R van der, Mulder WJM, Netea MG. Trained immunity: Target for prophylaxis and therapy. Cell Host Microbe. 2023; 31(11):1776–1791.

8. Nemeth J, Olson GS, Rothchild AC, et al. Contained Mycobacterium tuberculosis infection induces concomitant and heterologous protection. PLoS pathogens [Internet]. 2020; 16(7):e1008655–25. Available from: 10.1371/journal.ppat.1008655

9. Mai D, Jahn A, Murray T, et al. Mycobacterial exposure remodels alveolar macrophages and the early innate response to Mycobacterium tuberculosis infection. Biorxiv. 2022; :2022.09.19.507309.

10. Kusejko K, Günthard HF, Olson GS, et al. Diagnosis of latent tuberculosis infection is associated with reduced HIV viral load and lower risk for opportunistic infections in people living with HIV. Rowland-Jones SL, editor. PLOS Biology. 2020; 18(12):e3000963–16.

11. Tepekule B, Kusejko K, Zeeb M, et al. Impact of Latent Tuberculosis on Diabetes. J Infect Dis. 2022;.

12. Scherrer AU, Traytel A, Braun DL, et al. Cohort Profile Update: The Swiss HIV Cohort Study (SHCS). Int J Epidemiology. 2021; 51(1):33–34j.

13. Bray NL, Pimentel H, Melsted P, Pachter L. Near-optimal probabilistic RNA-seq quantification. Nat Biotechnol. 2016; 34(5):525–527.

14. Love MI, Huber W, Anders S. Moderated estimation of fold change and dispersion for RNA-seq data with DESeq2. Genome Biol. 2014; 15(12):550.

15. Ritchie ME, Phipson B, Wu D, et al. limma powers differential expression analyses for RNA-sequencing and microarray studies. Nucleic Acids Res. 2015; 43(7):e47–e47.

16. Kaforou M, Wright VJ, Oni T, et al. Detection of Tuberculosis in HIV-Infected and - Uninfected African Adults Using Whole Blood RNA Expression Signatures: A Case-Control Study. Cattamanchi A, editor. PLoS medicine. 2013; 10(10):e1001538–17.

17. Sweeney TE, Braviak L, Tato CM, Khatri P. Genome-wide expression for diagnosis of pulmonary tuberculosis: a multicohort analysis. Lancet Respir Med. 2016; 4(3):213–224.

18. Rotger M, Dang KK, Fellay J, et al. Genome-Wide mRNA Expression Correlates of Viral Control in CD4+ T-Cells from HIV-1-Infected Individuals. Plos Pathog. 2010; 6(2):e1000781.

19. Toossi Z, MayanjaLKizza H, Hirsch CS, et al. Impact of tuberculosis (TB) on HIVL1 activity in dually infected patients. Clin Exp Immunol. 2001; 123(2):233–238.

20. Nemeth J, Winkler H-M, Boeck L, et al. Specific cytokine patterns of pulmonary tuberculosis in Central Africa. Clinical immunology (Orlando, Fla). 2011; 138(1):50–59.

21. Verbon, Juffermans, Deventer V, Speelman, Deutekom V, Poll VD. Serum concentrations of cytokines in patients with active tuberculosis (TB) and after treatment. Clin Exp Immunol. 1999; 115(1):110–113.

22. Gupte AN, Kumar P, Araújo-Pereira M, et al. Baseline IL-6 is a biomarker for unfavorable tuberculosis treatment outcomes: a multi-site discovery and validation study. Eur Respir J. 2022; 59(4):2100905.

23. Martinez AN, Mehra S, Kaushal D. Role of Interleukin 6 in Innate Immunity to Mycobacterium tuberculosis Infection. J Infect Dis. 2013; 207(8):1253–1261.

24. Temu TM, Polyak SJ, Wanjalla CN, et al. Latent tuberculosis is associated with heightened levels of pro-and anti-inflammatory cytokines among Kenyan men and women living with HIV on long-term antiretroviral therapy. AIDS. 2023; 37(7):1065–1075.

25. Tabb B, Morcock DR, Trubey CM, et al. Reduced Inflammation and Lymphoid Tissue Immunopathology in Rhesus Macaques Receiving Anti–Tumor Necrosis Factor Treatment During Primary Simian Immunodeficiency Virus Infection. J Infect Dis. 2013; 207(6):880–892.

26. Vaidya SA, Korner C, Sirignano MN, et al. Tumor Necrosis Factor α Is Associated With Viral Control and Early Disease Progression in Patients With HIV Type 1 Infection. J Infect Dis. 2014; 210(7):1042–1046.

27. Plessner HL, Lin PL, Kohno T, et al. Neutralization of tumor necrosis factor (TNF) by antibody but not TNF receptor fusion molecule exacerbates chronic murine tuberculosis. The Journal of infectious diseases. 2007; 195(11):1643–1650.

28. Botha T, Ryffel B. Reactivation of latent tuberculosis infection in TNF-deficient mice. 2003; 171(6):3110–3118.

29. Clay H, Volkman HE, Ramakrishnan L. Tumor necrosis factor signaling mediates resistance to mycobacteria by inhibiting bacterial growth and macrophage death. Immunity. 2008; 29(2):283–294.

30. Díez-Fuertes F, Torre-Tarazona Hedl, Calonge E, et al. Transcriptome Sequencing of Peripheral Blood Mononuclear Cells from Elite Controller-Long Term Non Progressors. Sci Rep-uk. 2019; 9(1):14265.

31. Bates M, Mudenda V, Shibemba A, et al. Burden of tuberculosis at post mortem in inpatients at a tertiary referral centre in sub-Saharan Africa: a prospective descriptive autopsy study. The Lancet Infectious diseases. 2015; 15(5):544–551.

32. Uhlén M, Fagerberg L, Hallström BM, et al. Proteomics. Tissue-based map of the human proteome. Sci (N York, NY). 2015; 347(6220):1260419.

