## Supplementary Material for "*Mycobacterium tuberculosis* infection associated immune perturbations correlate with antiretroviral immunity"

**Supplementary Material for the manuscript entitled "*Mycobacterium tuberculosis* infection associated immune perturbations correlate with antiretroviral immunity"**

Burcu Tepekule^*,1,2,3^, Lisa Jörimann^*,1,2^, Corinne D. Schenkel^1,2^, Lennart Opitz^4^, Jasmin Tschumi^1,2^, Rebekka Wolfensberger^1^, Kathrin Neumann^1,2^, Katharina Kusejko^1^, Marius Zeeb^1,2^, Lucas Boeck^5^, Marisa Kälin^1^, Julia Notter^6^, Hansjakob Furrer^7^, Matthias Hoffmann^8^, Hans H. Hirsch^9,10,11^, Alexandra Calmy^12^, Matthias Cavassini^13^, Niklaus D. Labhardt^14,15^, Enos Bernasconi^12,16,17^, Karin J Metzner^1,2^, Dominique L. Braun^1,2^, Huldrych F. Günthard^1,2^, Roger D. Kouyos^1,2^, Fergal Duffy^18^, Johannes Nemeth^1^ and the Swiss HIV Cohort Study.

* These authors contributed equally

^1^ Department of Infectious Diseases and Hospital Epidemiology, University Hospital Zurich, Zurich, Switzerland.

^2^ Institute of Medical Virology, University of Zurich, Zurich, Switzerland.

^3^ Department of Ecology and Evolutionary Biology, Princeton University, Princeton, USA.

^4^ Functional Genomics Center Zurich, Swiss Federal Institute of Technology and University of Zurich, Zurich, Switzerland.

^5^ Department of Biomedicine, University of Basel, Switzerland

^6^ Division of Infectious Diseases and Hospital Epidemiology, Cantonal Hospital St Gallen, St. Gallen, Switzerland.

^7^ Department of Infectious Diseases, Inselspital, Bern University Hospital, University of Bern, Bern, Switzerland.

^8^ Division of Infectious Diseases and Hospital Epidemiology, Cantonal Hospital Olten, Olten, Switzerland

^9^ Division of Infectious Diseases and Hospital Epidemiology, University Hospital Basel, Basel, Switzerland.

^10^ Clinical Virology, Laboratory Medicine, University Hospital Basel, Basel, Switzerland.

^11^ Department Biomedicine, Transplantation and Clinical Virology, University of Basel, Basel, Switzerland.

^12^ Division of Infectious Diseases, University Hospital Geneva, University of Geneva, Geneva, Switzerland.

^13^ Division of Infectious Diseases, University Hospital Lausanne, University of Lausanne, Lausanne, Switzerland.

^14^ Division Clinical Epidemiology, Department of Clinical Research, University Hospital Basel, Basel, Switzerland.

^15^ University of Basel, Basel, Switzerland.

^16^ Division of Infectious Diseases, Ente Ospedaliero Cantonale, Lugano, Switzerland.

^17^ University of Geneva and University of Southern Switzerland, Lugano, Switzerland.

^18^ Seattle Children's Research Institute, Seattle, Washington, USA

### **Corresponding author**

Dr. med. Johannes Nemeth

Senior physician

Department of Infectious Diseases and Hospital Epidemiology

University Hospital Zurich

Rämistrasse 100

8091 Zürich

Switzerland

+41 44 255 33 22

**Keywords :** Tuberculosis, HIV, HIV-TB co-infection, immune system, transcriptome

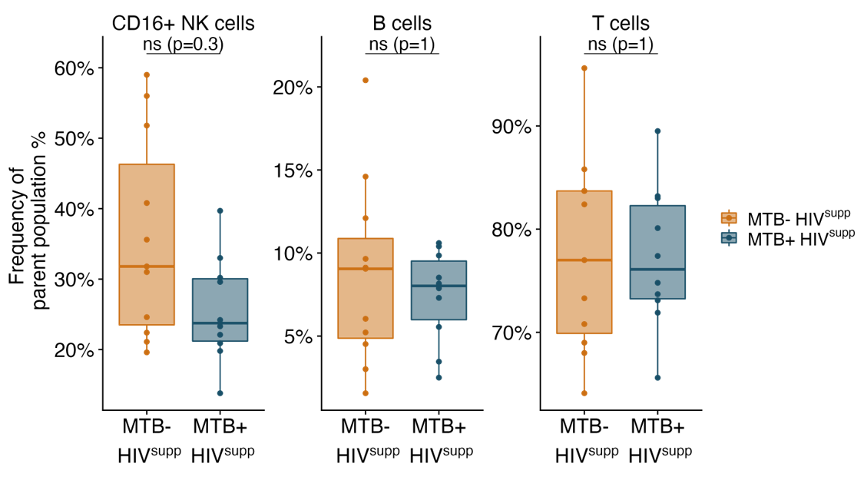

**Fig. S1a. Flow cytometry analysis for HIV^supp^ individuals.** Frequency of CD16+ NK cells, B cells and T cells in MTB+ HIV^supp^ (blue) and MTB- HIV^supp^ (orange) after flow cytometry analysis. Shown are boxplots with median and standard deviation. Wilcoxon test is shown for each cell type (CD16+ NK cells: p=0.3, B cells: p=1, T cells: p=1). CD16+ NK cell (MTB+ HIVsupp: 1.62 (1.14, 1.97), MTB- HIVsupp: 2.13 (1.05, 4.16), median (IQR), p=0.3), B cell (MTB+ HIVsupp: 4.55 (3.32, 5.46), MTB- HIVsupp: 4.42 (3.22, 6.97), median (IQR), p=1) or T cell (MTB+ HIVsupp: 50.5 (36.6, 59.8), MTB- HIVsupp: 55.7 (48.4, 58.4), median (IQR), p=1)

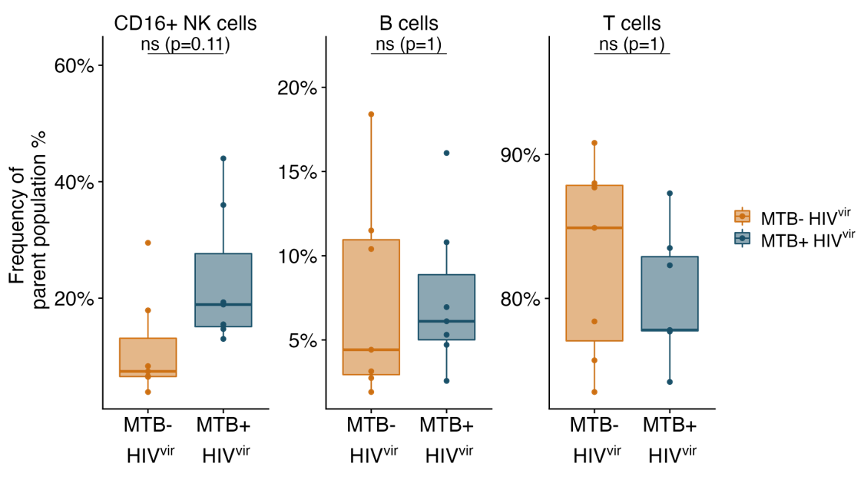

**Fig. S1b. Flow cytometry analysis for HIV^vir^ individuals.** Frequency of CD16+ NK cells, B cells and T cells in MTB+ HIV^vir^ (blue) and MTB- HIV^vir^ (orange) after flow cytometry analysis. Shown are boxplots with median and standard deviation. Wilcoxon test is shown for each cell type (CD16+ NK cells: p=0.11, B cells: p=1, T cells: p=1).

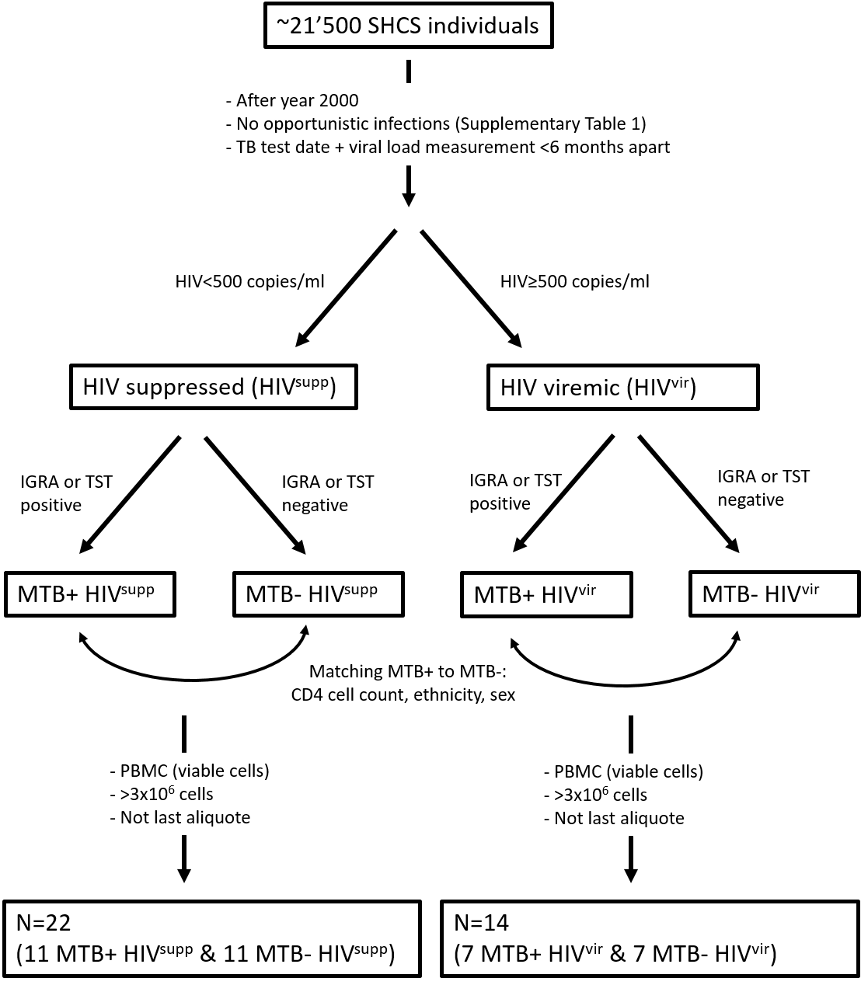

**Fig. S2.** **Flow chart of selection criteria for PWH.** Shown are number of individuals after each selection step and the corresponding selection criteria.

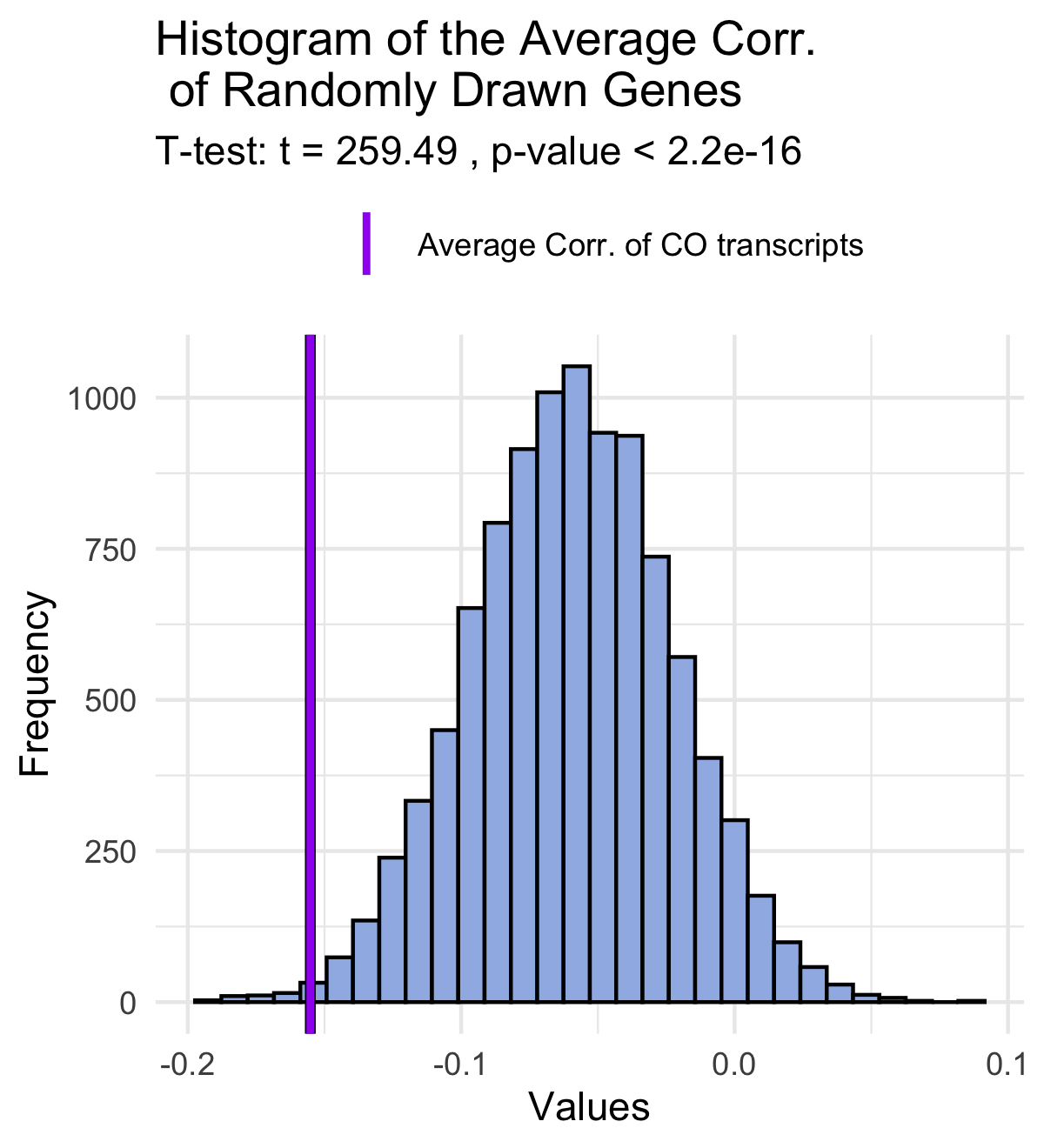

**Figure S3. b)** Histogram of the average correlation between HIV-1 viral load and the expression levels for each randomized set generated during the permutation test.

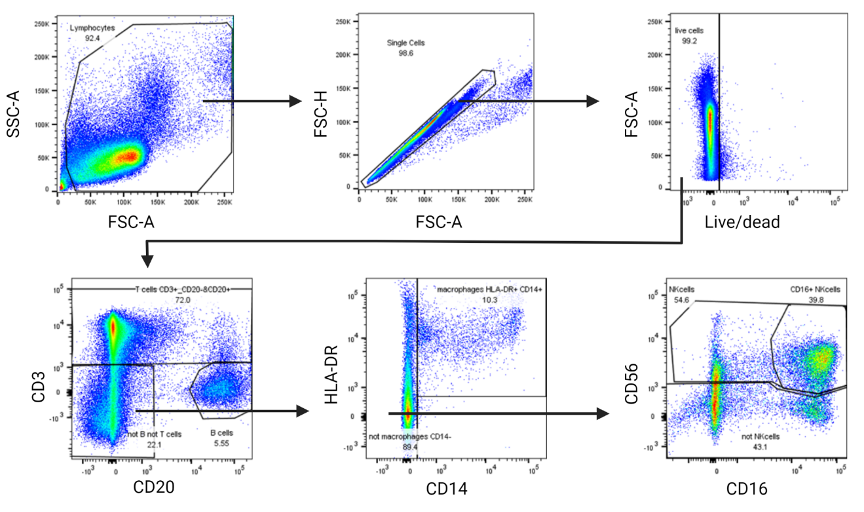

**Fig. S3. Gating strategy for flow cytometry analysis.** PBMCs from PWH were obtained and live cells were gated on CD3 and CD20. CD3-/CD20- cells were further gated on HLA-DR and CD14 and CD14- cells were further gated on CD56 and CD16.

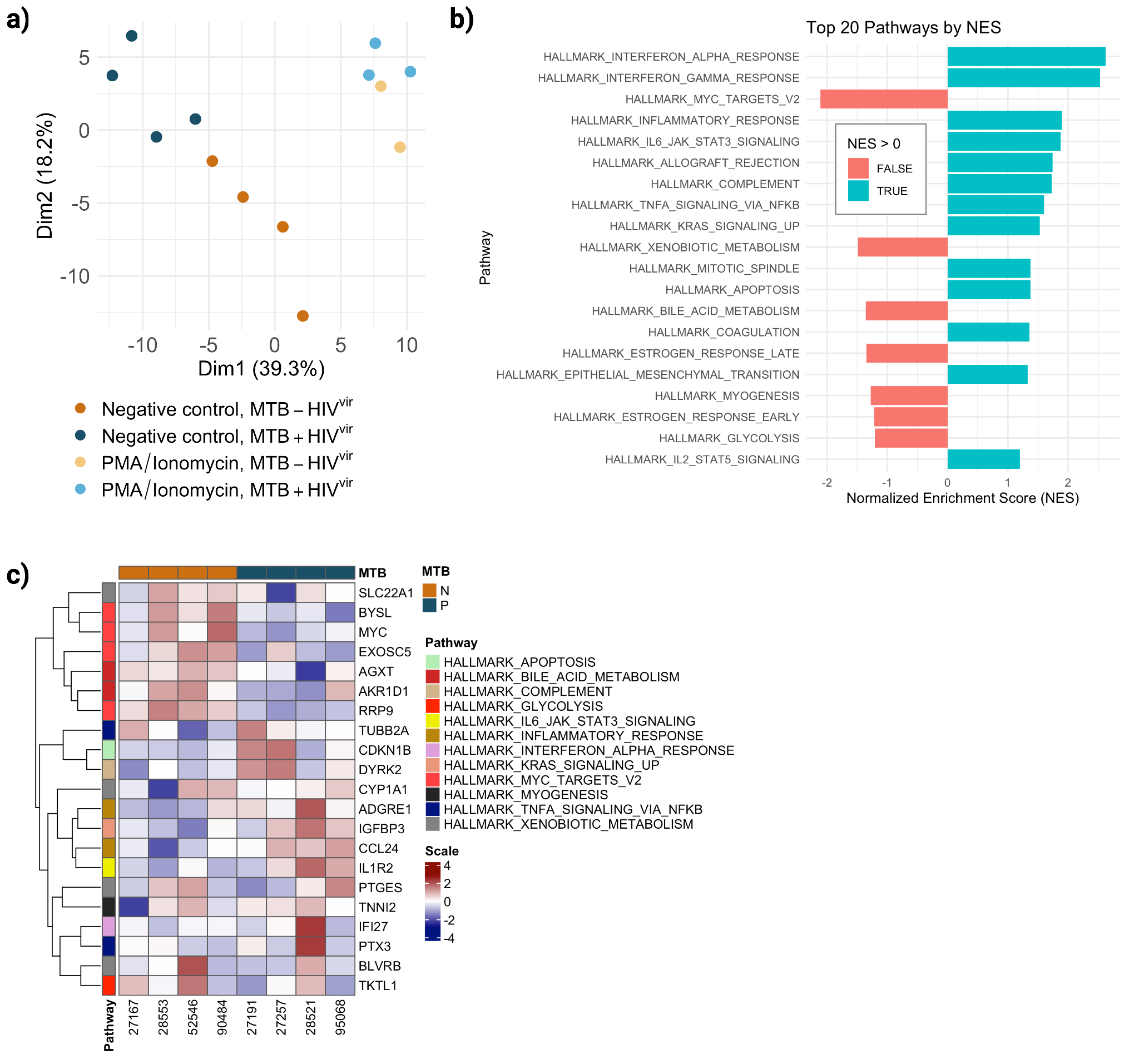

**Fig. S4. MTB Perturbations in viremic PWH, including only 4 matched pairs for HIV^vir^. a)** Dimensionality reduction and clustering analysis for HIV-viremic individuals with and without MTB infection (MTB+ HIV^vir^ and MTB- HIV^vir^). **b)** First 20 pathways with the highest normalized enrichment score (NES) based on the gene-set enrichment analysis (GSEA). **c)** Heatmap displaying the leading edge genes of these pathways filtered by p-value <0.01.

**Table S1. List of carried over transcripts.** List of carried over transcripts with their corresponding names, log fold change values, p values, and correlation with the viral load.

| gene_name | log2_HIVP | pValue_HIVP | log2_HIVN | pValue_HIVN | corr_FPKM |
| --- | --- | --- | --- | --- | --- |
| AMY2B | 1.498517 | 0.0054020076 | 0.05397872 | 0.77417492 | -0.757981518 |
| NEO1 | 2.097385 | 0.0003659422 | 0.25259358 | 0.48651145 | -0.695375674 |
| TUBB2A | 1.356838 | 0.0038829315 | 0.25562511 | 0.52933326 | -0.693195713 |
| AC008581.2 | 3.607487 | 0.0016574965 | 0.93509091 | 0.13561844 | -0.618131888 |
| CEACAM21 | 1.556890 | 0.0005316478 | 0.02100271 | 0.94718182 | -0.600885231 |
| TUBB2B | 4.283380 | 0.0001594871 | 0.20081191 | 0.79685463 | -0.588355761 |
| WWC2 | 1.552436 | 0.0036873349 | 0.31473831 | 0.11974292 | -0.468554078 |
| AC010616.1 | 2.198713 | 0.0014058191 | 0.22076107 | 0.73842338 | -0.352098246 |
| GNLY | 2.202992 | 0.0012511819 | 0.08453818 | 0.87513189 | -0.339678519 |
| BX664615.2 | 2.844257 | 0.0002237700 | 0.37803702 | 0.39299079 | -0.337670751 |
| IGFBP3 | 1.074670 | 0.0037012894 | 0.16686169 | 0.78135045 | -0.167691499 |
| AC020915.4 | 5.249703 | 0.0001215270 | 0.30825722 | 0.61147559 | -0.158615727 |
| SNED1 | 1.550061 | 0.0097260470 | 0.57065683 | 0.08013222 | -0.152394369 |
| CCBE1 | -1.300812 | 0.0030550573 | -0.02680895 | 0.93412474 | -0.145279671 |
| XAB2 | 1.204309 | 0.0032352300 | 0.80578373 | 0.05997778 | -0.139487287 |
| CDCA8 | 1.419941 | 0.0038365777 | 0.16759541 | 0.56798927 | -0.070498826 |
| IGKV1-12 | -1.733855 | 0.0016667139 | -0.38806199 | 0.58113219 | -0.020101140 |
| SIGLEC6 | -1.430166 | 0.0012398728 | -0.03619391 | 0.90171577 | -0.007168388 |
| RERG | -1.500063 | 0.0056277682 | -0.70385719 | 0.06562413 | -0.006620515 |
| AC018695.7 | -1.325224 | 0.0083501188 | -0.29446126 | 0.38627728 | -0.003516010 |
| STAC2 | -1.013961 | 0.0065117858 | -0.12694852 | 0.69753877 | 0.016734316 |
| IGHV4-59 | -1.563449 | 0.0032762821 | -0.46801202 | 0.31071520 | 0.023350983 |
| AP000295.1 | 7.895425 | 0.0058325832 | 1.31325802 | 0.18259731 | 0.032797403 |
| IGKV3-7 | -2.051104 | 0.0029566524 | -0.81677588 | 0.43345342 | 0.097574788 |
| IGHV1-69D | 1.561447 | 0.0048548079 | 0.43449272 | 0.59827189 | 0.169790697 |
| HS3ST3B1 | 2.105495 | 0.0077163204 | 0.63054452 | 0.19011784 | 0.237958915 |
| AKR1D1 | -1.175365 | 0.0070091463 | -0.37013928 | 0.27652881 | 0.277186669 |
| CYP1A1 | -2.043803 | 0.0049102523 | -0.30249073 | 0.66090913 | 0.277646563 |
| FAM156A | -1.516044 | 0.0073136239 | -0.46891545 | 0.17543333 | 0.277827248 |
| PDE4C | -1.035886 | 0.0013943918 | -0.15551315 | 0.59732671 | 0.382744588 |
| RAB34 | -1.522200 | 0.0012275430 | -0.47922421 | 0.24850641 | 0.423963744 |
| SFTPB | -1.090287 | 0.0051743503 | -0.14841951 | 0.65932146 | 0.464057520 |

**Table S2. List of opportunistic infections that were excluded for selection of PWH**

Aids defining disease not specified

Bacterial pneumonia, recurrent

Cytomegalovirus (CMV) - retinitis

Cytomegalovirus (CMV) disease, other

Candidiasis of trachea, bronchi or lungs

Candidiasis, oesophagial

Carcinoma, cervical, invasive

Coccidioidomycosis disseminated

Cryptococcal meningitis

Cryptococcosis, other disseminated

Cryptosporidiosis, Diarrhea > 1 month

Encephalopathy, HIV-related

Herpes simplex disease, visceral

Histoplasmosis disseminated

Intracerebral lesions, indeterminate

Isosporiasis , Diarrhoe > 1 month

Kaposi sarcoma

Mycobacterium avium - intracellulare, disseminated

Mycobacterium genavense disease

Mycobacterium kansasii disease

Mycobacterium avium complex or kansasii

Mycobacterium other species disseminated or extrapulmonary

Non-Hodgkin's lymphoma

Pediatric category C disease

Pneumocystis disease, extrapulmonary

Pneumocystis pneumonia

Salmonella septicemia, recurrent

Toxoplasmosis disseminated

Toxoplasmosis, cerebral

Tuberculosis pulmonary

Wasting Syndrome, AIDS-defining
